# Pangenome graph layout by Path-Guided Stochastic Gradient Descent

**DOI:** 10.1101/2023.09.22.558964

**Authors:** Simon Heumos, Andrea Guarracino, Jan-Niklas M. Schmelzle, Jiajie Li, Zhiru Zhang, Jörg Hagmann, Sven Nahnsen, Pjotr Prins, Erik Garrison

## Abstract

**Motivation:** The increasing availability of complete genomes demands for models to study genomic variability within entire populations. Pangenome graphs capture the full genomic similarity and diversity between multiple genomes. In order to understand them, we need to see them. For visualization, we need a human readable graph layout: A graph embedding in low (e.g. two) dimensional depictions. Due to a pangenome graph’s potential excessive size, this is a significant challenge.

**Results:** In response, we introduce a novel graph layout algorithm: the Path-Guided Stochastic Gradient Descent (PG-SGD). PG-SGD uses the genomes, represented in the pangenome graph as paths, as an embedded positional system to sample genomic distances between pairs of nodes. This avoids the quadratic cost seen in previous versions of graph drawing by Stochastic Gradient Descent (SGD). We show that our implementation efficiently computes the low dimensional layouts of gigabase-scale pangenome graphs, unveiling their biological features.

**Availability:** We integrated PG-SGD in *ODGI* which is released as free software under the MIT open source license. Source code is available at https://github.com/pangenome/odgi.

## 1 Introduction

Reference genomes are widely used in genomics, serving as a foundation for a variety of analyses, including gene annotation, read mapping, and variant detection (Singh *et al*., 2022). However, this linear model is becoming obsolete given the accessibility to hundreds or even thousands of high-quality genomes. A single genome can not fully represent the genetic diversity of any species, resulting in reference bias (Ballouz *et al*., 2019). In contrast, a pangenome models the entire set of genomic elements of a given population (Tettelin *et al*., 2008; Computational Pan-Genomics Consortium, 2018; Eizenga *et al*., 2020; Sherman and Salzberg, 2020). Pangenomes can be represented as a sequence graph incorporating sequences as nodes and their relationships as edges (Hein, 1989). In the variation graph model (Garrison *et al*., 2018), genomes are encoded as paths traversing the nodes in the graph.

A graph layout is the arrangement of nodes and edges in an *N* - dimensional space. Graph layout algorithms aim to find optimal node coordinates in order to minimize overlapping nodes or edges, reduce edge crossings, and promote an intuitive understanding of the graph. One popular approach is force-directed graph drawing (Cheong and Si, 2022) which produces aesthetic layouts. This is prone to get stuck in local minima, but stochastic gradient descent (SGD) implementations alleviate such a problem (Zheng *et al*., 2019). SGD uses the gradient of its individual terms to approximate the gradient of a sum of functions.

A *pangenome* graph layout can provide a human-readable visualization of genetic variation between multiple genomes. However, Zheng *et al*. (2019)’s algorithm has a quadratic up front cost in the number of nodes to find pairwise distances to guide the layout, making it impossible to apply to pangenome graphs with millions of nodes. Also, existing graph layout approaches ignore the biological information inherent in pangenome graphs.

In practice, multidimensional scaling (MDS) is applied to minimize the difference between the visual distance and theoretical graph distance. This can be accomplished by using pairwise node distances to minimize an energy function. Since pangenome graphs represent genomes as paths in the graph, a reasonable distance metric would be the nucleotide distance between a pair of nodes traversed by the same path. Such path sampling would overcome the quadratic costs of previous versions of graph drawing by SGD.

Typically, force-directed layouts are hard to compute (Wang *et al*., 2014), but the lock-free HOGWILD! method offers a highly parallelizable and thus scalable SGD approach that can be applied when the optimization problem is sparse (Recht *et al*., 2011).

Here, we present a new pangenome graph layout algorithm which applies a path-guided stochastic gradient descent (PG-SGD) to use the paths as an embedded positional system to find distances between nodes, moving pairs of nodes in parallel with a modified HOGWILD! strategy. The algorithm computes the pangenome graph layout that best reflects the nucleotide sequences in the graph. To our knowledge, no algorithm takes into account such biological information to compute the graph layouts. PG-SGD can be extended in any number of dimensions. In the ODGI toolkit (Guarracino *et al*., 2022), we provide implementations for 1-dimensional (1D) and 2-dimensional (2D) layouts. These algorithms have already been successfully applied to construct and visualize large-scale pangenome graphs of the Human Pangenome Reference Consortium (HPRC) (Liao *et al*., 2023; Guarracino *et al*., 2023).

## 2 Algorithm

While PG-SGD is inspired by Zheng *et al*. (2019), we designed the algorithm to work on the variation graph model (Definition 2.1).

### Definition 2.1.

Variation graphs are a mathematical formalism to represent pangenome graphs (Garrison, 2019). In the variation graph *𝒢* = (*𝒱, ℰ, 𝒫*), nodes (or vertices) *𝒱* = *v*_1_ … *v*| *𝒱*| contain nucleotide sequences. Each node *v*_*i*_ has a unique identifier *i* and an implicit reverse complement 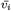. The node strand *o* represents the node orientation. Edges *ℰ* = *e*_1_ … *e* _| *ℰ* |_ connect ordered pairs of node strands (*e*_*i*_ = (*o*_*a*_, *o*_*b*_)), defining the graph topology. Paths *𝒫* = *p*_1_ … *p* _|*P*|_ are series of connected steps *s*_*i*_ that refer to node strands in the graph (*p*_*i*_ = *s*_1_ … *s* _|*pi*|_); the paths represent the genomes embedded in the graph.

We report PG-SGD’s pseudocode in Algorithm 1 and its schematic in Figure 1. In brief, the algorithm moves one pair of nodes (*v*_*i*_, *v*_*j*_) at a time, minimizing the difference between the layout distance *ld*_*ij*_ of the two nodes and the nucleotide distance *nd*_*ij*_ of the same nodes as calculated along a path that traverses them. In the 2D layouts, nodes have two ends. When moving a pair of nodes, we actually move one end of each node.

**Fig. 1.**
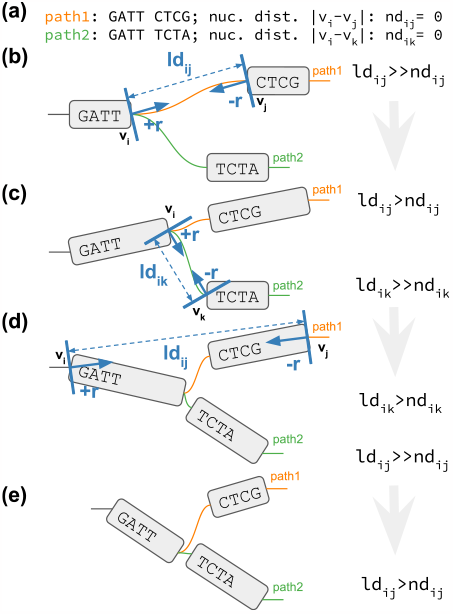
2D PG-SGD update operation sketches. **(a)** The path information of the graph. *path1* and *path2* both visit the same first node. Then their sequence diverges and they visit distinct nodes. **(b-e)** *v*_*i*_/*v*_*j*_ or *v*_*i*_/*v*_*k*_ is the current pair of nodes to update. *ld*_*ij*_ /*ld*_*ik*_ is the current layout distance. *r, ™r* is the current size of the update. **(b)** Initial graph layout highlighting the future update of the two nodes of *path1*. **(c)** The graph layout after the first update. The nodes appear longer now, because we updated at the end of the nodes. Highlighted is the future update of the two nodes of *path2*. **(d)** The graph layout after the second update. Highlighted is the future update of the two nodes of *path1*. **(e)** Final graph layout after three updates using the 2D PG-SGD.

For clarification, an example is given in Figure 1. *v*_*i*_ is the node associated with the step *s*_*i*_ sampled uniformly from all the steps in *P. v*_*j*_ is the node associated with the step *s*_*j*_ sampled from the same path of *s*_*i*_ by drawing a uniform or a Zipfian distribution (Zipf, 1932). The difference between *Heumos, Guarracino et al*.*nd*_*ij*_ and *ld*_*ij*_ guides the update of the node coordinates in the layout. The magnitude *r* of the update depends on the learning rate *µ*. The number of iterations steers the annealing step size *η* which determines the learning rate *µ*. A large *η* in the first iterations leads to a globally linear (in 1D) or planar (in 2D) layout. By decreasing *η*, the layout adjustments become more localized, ensuring that the nodes are positioned to best reflect the nucleotide distances in the paths (i.e., in the genomes).

### Algorithm 1

Pseudocode of PG-SGD in 1D.

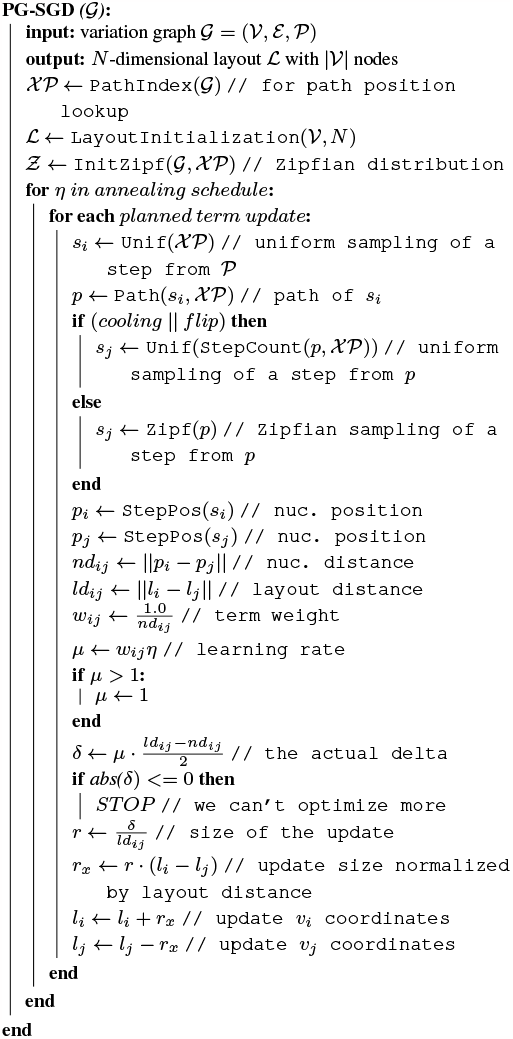

Originating from empirical inspection of word frequency tables, Zipf’s law states that a word with rank *n* occurs 1*/n* times as the most frequent one. This law is modeled by the Zipf distribution.Sampling *s*_*j*_ from a Zipf distribution fixed in the *s*_*i*_’s path position space increases the possibility to draw a nucleotide position close to *s*_*i*_. So there is a high chance to use small nucleotide distances *nd*_*ij*_ to refine the layout of nodes comprising a few base pairs. The Zipf distribution is also long-tailed, with many occurrences of low frequency events. However, extremely long-range correlations might not be captured sufficiently, resulting in collapsed layouts for structures that are otherwise linear. To provide “main” — 2023/10/17 — 15:54 — page 3 — #3 *Pangenome graph layout by Path-Guided Stochastic Gradient Descent* **3** balance between global and local layout updates, in half of the updates (*flip* flag in Algorithm 1), the *s*_*j*_ is sampled uniformly instead from a Zipf distribution, with uniform sampling being more favorable for global updates. Furthermore, to enhance local linearity (in 1D) or planarity (in 2D) of the graph layout, a *cooling* phase skews the Zipfian distribution after half of iterations have been completed. This increases the likelihood of sampling smaller nucleotide distances for the layout updates.

## 3 Implementation

We implemented PG-SGD in ODGI (Guarracino *et al*., 2022): the 1D version can be found in *odgi sort* and the 2D version in *odgi layout*. To efficiently retrieve path nucleotide positions, we implemented a path index. This index is a strict subset of the XG index (Garrison *et al*., 2018) where we avoid to use succinct SDSL data structures (Gog *et al*., 2014). Instead, we rely on bit-compressed integer vectors, enabling efficient retrieval of path nucleotide positions to quickly compute nucleotide distances without having to store all pairwise distances between nodes in memory. This approach ensures to scale on large pangenome graphs representing thousands of whole genomes.

Graph layout initialization can significantly influence the quality of the final layout. In the 1D implementation, by default, nodes are placed in the same order as they appear in the input graph, although we also provide support for random layout initialization. In 2D, we offer several layout initialization techniques. One approach places nodes in the first layout dimension according to their order in the input graph, adding either uniform or Gaussian noise in the second dimension. Another strategy arranges nodes along a Hilbert curve, an approach that often favors the creation of planar final layouts. We also support fixing node positions to keep nodes in the same order as they are in a selected path, such as a reference genome. This feature allows us to build reference-focused graph layouts (Figure S1d).

Our implementation is multithreaded and uses shared memory for storing the layout in a vector, according to the HOGWILD! strategy (Recht *et al*., 2011). Threads perform layout updates without any locking for additional speed up. This approach is feasible since pangenome graphs are typically sparse (Guarracino *et al*., 2022), with low average node degree. As a result, the updates only modify small parts of the entire layout. While the HOGWILD! SGD algorithm writes the layout updates to a shared non-atomic double vector, PG-SGD stores node coordinates in a vector of atomic doubles. This vector prevents any potential memory overwrites. Our tests revealed basically no performance loss with respect to the non-atomic counterpart.

## 4 Results

### 4.1 Performance

We apply the 2D PG-SGD to the human pangenome (Liao *et al*., 2023) from the Human Pangenome Reference Consortium (HPRC) to show the scalability of the algorithm. Experiments were conducted on a cluster with 24 `Regular` nodes (32 cores / 64 threads with two AMD EPYC 7343 processors with 512 GB RAM) and 4 `HighMem` nodes (64 cores / 128 threads with two AMD EPYC 7513 processors with 2048 GB RAM). We downloaded pangenome graphs for each autosome (24 in total) and for the mitochondrial DNA. Each graph represents 90 whole human haplotypes: 44 diploid individuals plus the GRCh38 (Schneider *et al*., 2017) and CHM13 (Nurk *et al*., 2021) haploid human references (see Supplementary Table S1 for graph statistics). When applied to these pangenome graphs using one `Regular` node for each calculation, 2D PG-SGD obtains the graph layouts in 50 minutes on average, with the highest run time observed being chromosome 16 (Supplementary Table S1). This is expected since chromosome 16 has one of the highest levels of segmentally duplicated sequence among the human autosomes (Martin *et al*., 2004). Repetitive sequences lead to graph nodes with a very high number of path steps, which are computationally expensive to work with (Guarracino *et al*., 2022). Memory consumption is 29.66 GB of RAM on average, with the memory peak again occurring with chromosome 16, due to the path index building phase. Given its scalability, we even applied PG-SGD to the full graph with all chromosomes together using a HighMem node (Supplementary Table S1). *BandageNG* (https://github.com/asl/BandageNG, last accessed Jul 2023), the current state-of-the-art for graph visualization, was not able to produce a layout within 7 days, hitting the wall clock time limit of the cluster. On average, PG-SGD is *∼*8X faster than BandageNG while using *∼*2X less memory.

### 4.2 Pangenome graph layouts reveal biological features

Graph visualization is essential for understanding pangenome graphs and the genome variation they represent. We show how 2D PG-SGD allows us gaining insight into biological data by looking at the graph layout structure. In Figure 2a, the chromosomes of the HPRC graph show the large scale structural variations in the centromeres. Focusing on the major histocompatibility complex (MHC) of chromosome 6 (Figure 2b), the 2D layout reveals the positions and diversity of all MHC genes (Figure 2c). In Figure 2d the C4A and C4B genes are highlighted. Complementary, we provide various 1D visualizations in Supplementary Figure S1.

**Fig. 2.**
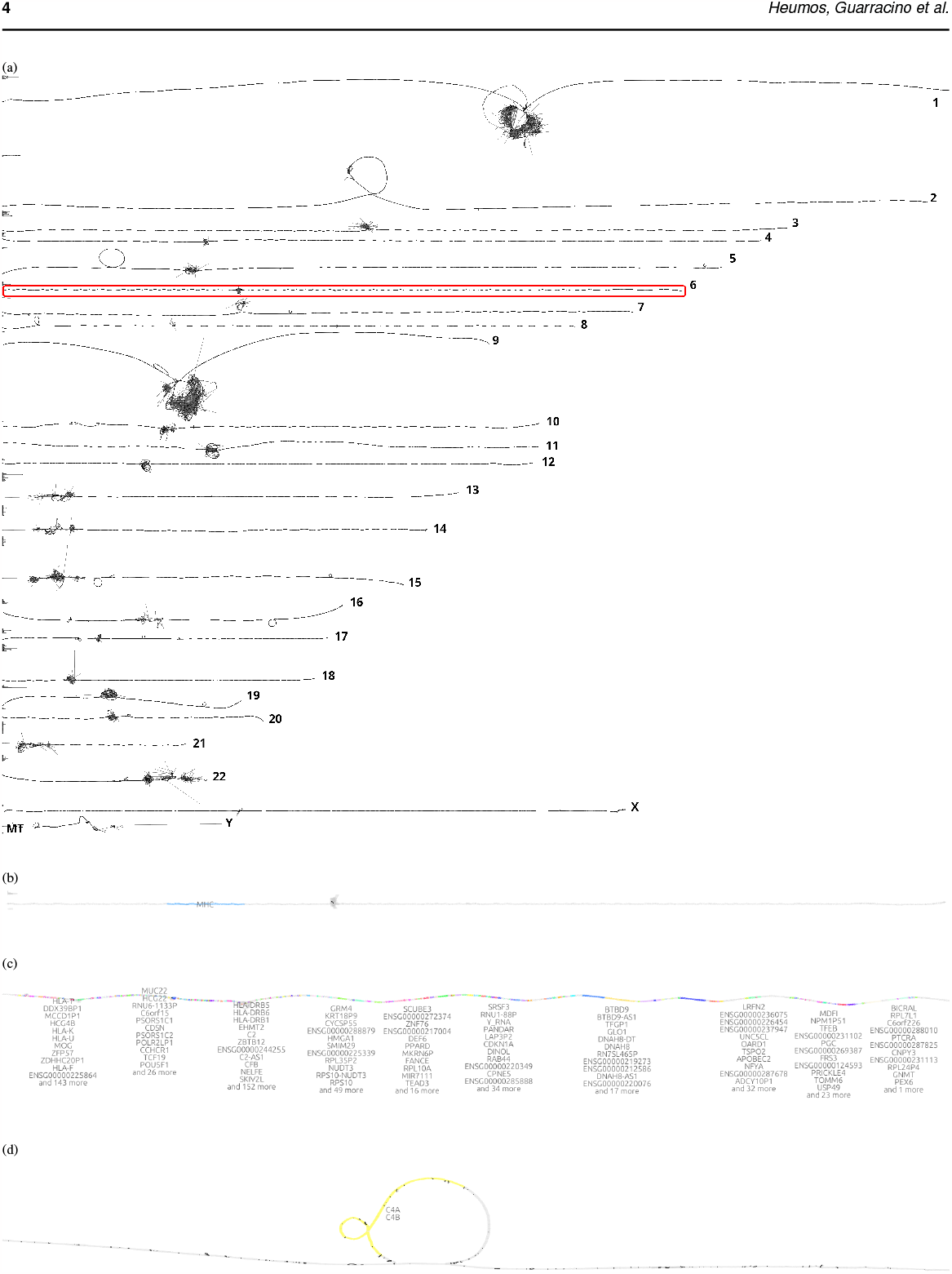
2D visualizations of all chromosomes of the Human Pangenome Reference Consortium (HPRC) 90 haplotypes pangenome graph, chromosome 6, the major histocompatibility complex (MHC), and the complement component 4 (C4). **(a)** *odgi draw* layout of the HPRC pangenome graph 90 haplotypes. Displayed are all 24 autosomes and the mitochondrial chromosome. A red rectangle highlights chromosome 6 which is shown in the subfigure below. **(b)** *gfaestus* screenshot of the chromosome 6 layout. Colored in blue is the MHC. The hairball in the middle is the centromere. The black structures in the centromere are edges. **(c)** *gfaestus* screenshot of the MHC. All MHC genes are color annotated and the names of the genes appear as a text overlay. **(d)** *gfaestus* screenshot of the region around C4, specifically color highlighting genes C4A and C4B. The black lines are the edges of the graph.

## 5 Discussion

We presented Path-Guided Stochastic Gradient Descent (PG-SGD), the first layout algorithm for pangenome graphs that leverages the biological information available within the genomes represented in the graph. Our implementation efficiently computes the layout of pangenome graphs representing thousands of whole genomes.

Graph visualization is key for understanding genome variations and the layouts produced by PG-SGD offer an unprecedented high-level perspective on pangenome variation. We implemented PG-SGD to generate layouts in 1D and 2D. These graph projections have already been employed in constructing and analyzing the first draft human pangenome reference (Liao *et al*., 2023), as well as in the discovery of heterologous recombination of human acrocentric chromosomes (Guarracino *et al*., 2023). Furthermore, they are applied in the creation and analysis of pangenome graphs for any species (Guarracino *et al*., 2022; Garrison *et al*., 2023). Of note, there still remains a gap in interactive and scalable solutions that merge layouts of large pangenome graphs with annotation. Our algorithm will underpin new pangenome graph browsers for studying graph layouts and the genome variation they represent (https://github.com/chfi/waragraph) last accessed Jul 2023).

The performance analysis shows that our 2D implementation outperforms *BandageNG* when handling large, complex pangenome graphs. While *BandageNG* was not able to deliver a layout of the whole HPRC graph within 1 week, our 2D PG-SGD calculated one within one day. There are some possible optimization approaches for future work to further improve the performance of PG-SGD, making it possible for interactive use. The data structure could be optimized to improve cache performance. Moreover, the high-degree of parallelism could be further exploited by using a GPU.

PG-SGD can be extended to any number of dimensions. It can be seen as a graph embedding algorithm that converts high-dimensional, sparse pangenome graphs into low-dimensional, dense, and continuous vector spaces, while preserving its biologically relevant information. This enables the application of machine learning algorithms that use the graph layout for variant detection and classification. Our future research involves leveraging these graph projections to detect structural variants and to identify and correct assembly errors. Moreover, we are considering extending the algorithm to RNA and protein sequences to support pantranscriptome graphs (Sibbesen *et al*., 2023) and panproteome graphs (Dabbaghie *et al*., 2023), respectively.

## Acknowledgments

We thank Matthias Seybold from the Quantitative Biology Center for maintaining the Core Facility Cluster.

## Software and data availability

Software versions, code, and links to data resources used to prepare this manuscript can be found at https://github.com/pangenome/sorting-paper. Animations of the algorithm are deposited at https://doi.org/10.5281/zenodo.8288999.

---

**Table S1:**
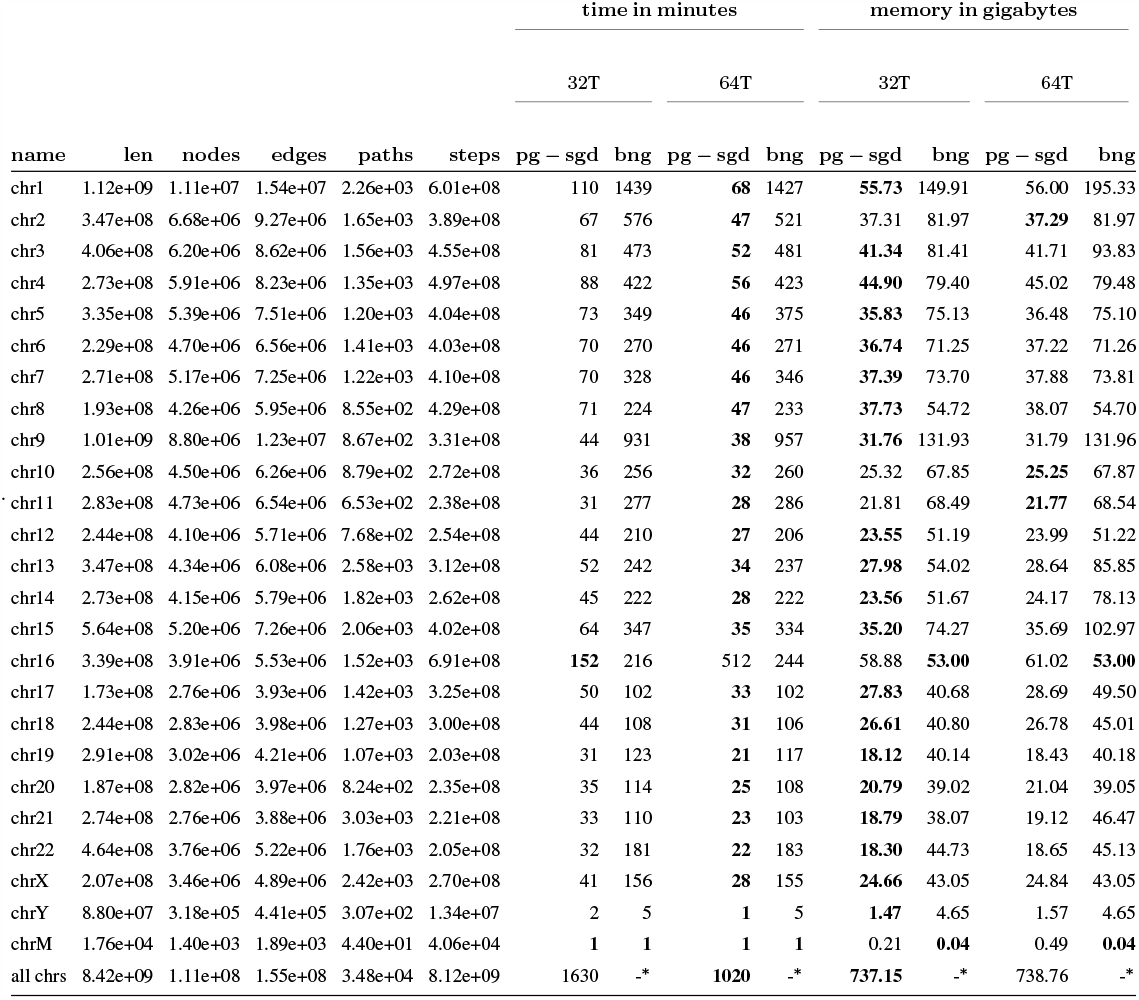
Performance evaluation of computing a 2D layout of all chromosomal HPRC pangenome graphs. From GFA to the actual layout. \**BandageNG* did not finish within the job wall clock time limit of 7 days. Therefore, no layout was produced. **32T:** Number of threads: 32. **64T:** Number of threads: 64.

## 6 Supplement

### 6.1 Supplementary data

#### 6.1.1 Performance evaluation

The results of the performance evaluation are given in Table S1.

#### 6.1.2 1D visualizations

The 1D PG-SGD algorithm creates a 1D layout of the nodes of the graph. Theoretically, it is possible that 2 nodes have the same 1D coordinate or overlap. But, in our 1D visualizations, we arrange the nodes from left to right. Therefore, we project the 1D coordinates into a 1D node order: We sort the final layout by graph component, graph position, and node rank.

**Fig. S1.**
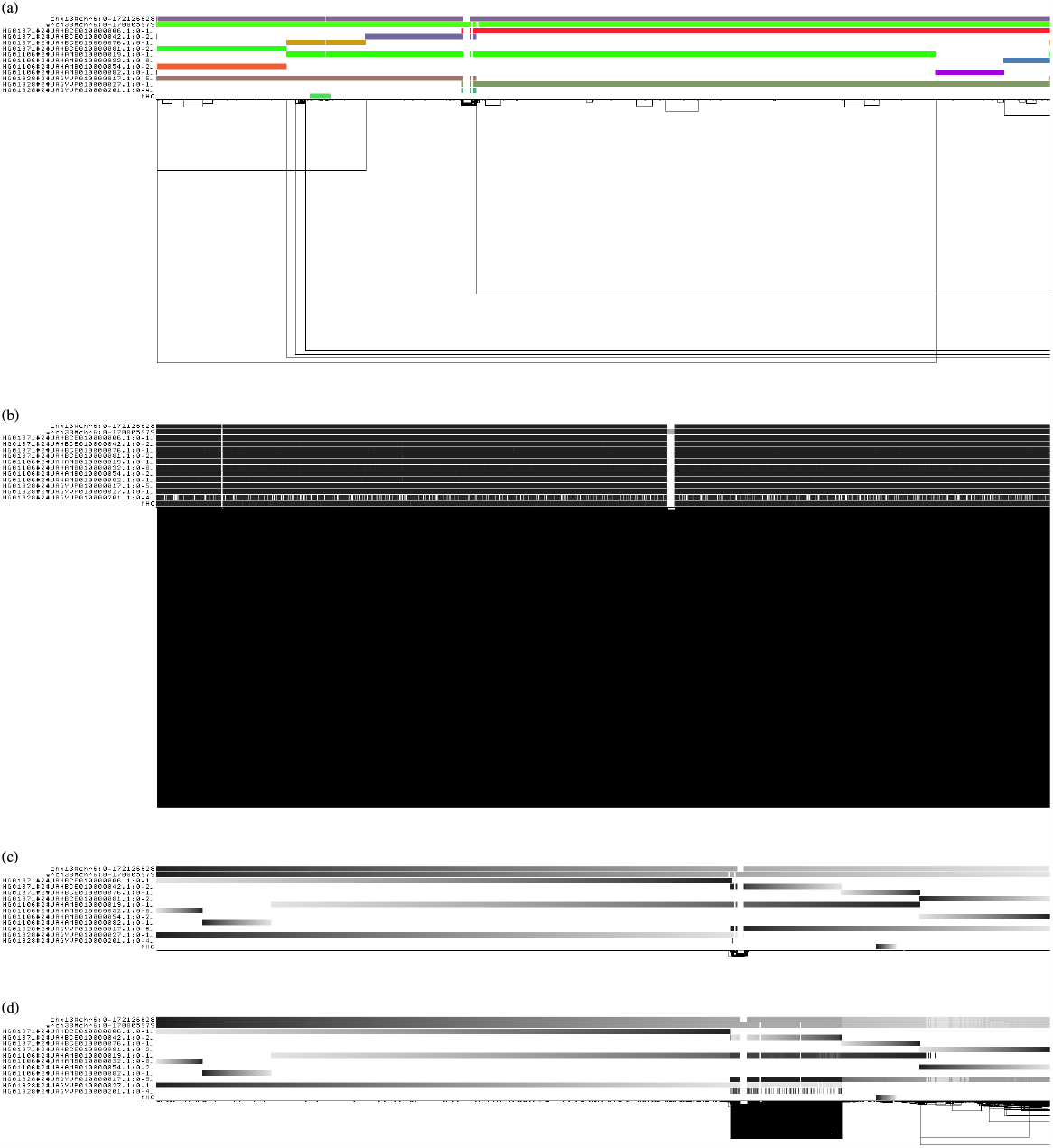
*odgi viz* 1D visualizations of a 5 haplotypes subgraph of the Human Pangenome Reference Consortium (HPRC) chromosome 6 pangenome graph. The major histocompatibility complex (MHC) sequence was injected as an extra path. Various node arrangements are shown. **a-d** A graphs nodes are arranged from left to right forming the pangenome sequence. The black lines under the paths are the links representing the topology. Path names are left. **(a)** *odgi viz* default modality: The colored bars are the paths versus pangenome sequence in a binary matrix. Shown is the subgraph extracted with *odgi extract*. No 1D layout algorithm was applied here. **b-d** *odgi viz* colored by path position. Light grey corresponds to the beginning of a path, black encodes the end of the path. **(b)** The nodes are arranged randomly in 1D. **(c)** The nodes are arranged applying the 1D PG-SGD algorithm. **(d)** The nodes are arranged applying a 1D reference-guided PG-SGD where the nodes of haplotype HG01071 are fixed and only all the othere ones are movable in 1D. Now all paths of this haplotype are arranged from their lowest to their highest nucleotide position. However,lot’s of longer links are now visible compared to the node ordering directly above. This indicates a node order that is globally not optimal.

## References

Ballouz, S. et al. (2019). Is it time to change the reference genome? Genome Biology, 20(1), 159.

Cheong, S.-H. and Si, Y.-W. (2022). Force-directed algorithms for schematic drawings and placement: A survey.

Computational Pan-Genomics Consortium (2018). Computational pan-genomics: status, promises and challenges. Briefings in Bioinformatics, 19(1), 118–135.

Dabbaghie, F. et al. (2023). PanPA: generation and alignment of panproteome graphs.

Eizenga, J. M. et al. (2020). Pangenome graphs. Annual Review of Genomics and Human Genetics, 21(1), 139–162.

Garrison, E. (2019). Graphical pangenomics.

Garrison, E. et al. (2018). Variation Graph Toolkit Improves Read Mapping by Representing Genetic Variation in the Reference. Nature Biotechnology, 36(9), 875–879.

Garrison, E. et al. (2023). Building pangenome graphs. bioRxiv.

Gog, S. et al. (2014). From theory to practice: Plug and play with succinct data structures. In 13th International Symposium on Experimental Algorithms, (SEA 2014), pages 326–337.

Guarracino, A. et al. (2022). ODGI: understanding pangenome graphs. Bioinformatics, 38(13), 3319–3326.

Guarracino, A. et al. (2023). Recombination between heterologous human acrocentric chromosomes. Nature, 617(7960), 335–343.

Hein, J. (1989). A new method that simultaneously aligns and reconstructs ancestral sequences for any number of homologous sequences, when the phylogeny is given. Molecular Biology and Evolution.

Liao, W.-W. et al. (2023). A draft human pangenome reference. Nature, 617(7960), 312–324.

Martin, J. et al. (2004). The sequence and analysis of duplication-rich human chromosome 16. Nature, 432(7020), 988–994.

Nurk, S. et al. (2021). The complete sequence of a human genome. BioRxiv.

Recht, B. et al./person-group>. (2011). Hogwild!: A Lock-Free Approach to Parallelizing Stochastic Gradient Descent. In J. Shawe-Taylor, R. Zemel, P. Bartlett, F. Pereira, and K. Q. Weinberger, editors, Advances in Neural Information Processing Systems, volume24. Curran Associates, Inc.

Schneider, V. A. et al. (2017). Evaluation of GRCh38 and de novo haploid genome assemblies demonstrates the enduring quality of the reference assembly. Genome Research, 27(5), 849–864.

Sherman, R. M. and Salzberg, S. L. (2020). Pan-genomics in the human genome era. Nature Reviews Genetics, 21(4), 243–254.

Sibbesen, J. A. et al. (2023). Haplotype-aware pantranscriptome analyses using spliced pangenome graphs. Nature Methods, 20(2), 239–247.

Singh, V. et al. (2022). From the reference human genome to human pangenome: Premise, promise and challenge. Frontiers in Genetics, 13.

Tettelin, H. et al. (2008). Comparative genomics: the bacterial pan-genome. Current Opinion in Microbiology, 11(5), 472–477.

Wang, L. et al. (2014). Research on Force-directed Algorithm Optimization Methods:. Shanghai, China.

Wick, R. R. et al. (2015). Bandage: interactive visualization of de novo genome assemblies. Bioinformatics, 31(20), 3350–3352.

Zheng, J. X. et al. (2019). Graph drawing by stochastic gradient descent. IEEE Transactions on Visualization and Computer Graphics, 25(9), 2738–2748.

Zipf, G. K. (1932). Selected Studies of the Principle of Relative Frequency in Language. Harvard University Press, Cambridge, MA and London, England.

